# Resource Competition Shapes the Response of Genetic Circuits

**DOI:** 10.1101/091306

**Authors:** Yili Qian, Hsin-Ho Huang, José I. Jimenéz, Domitilla Del Vecchio

**Author notes:** Correspondence: Address: Room 3-469, Department of Mechanical Engineering, Massachusetts Institute of Technology, 77 Massachusetts Avenue, Cambridge, MA02139, USA. These authors contributed equally to this work (Yili Qian, Hsin-Ho Huang).

## Abstract

A common approach to design genetic circuits is to compose gene expression cassettes together. While appealing, this modular approach is challenged by the fact that expression of each gene depends on the availability of transcriptional/translational resources, which is in turn determined by the presence of other genes in the circuit. This raises the question of how competition for resources by different genes affects a circuit’s behavior. Here, we create a library of genetic activation cascades in bacteria *E. coli*, where we explicitly tune the resource demand by each gene. We develop a general Hill-function-based model that incorporates resource competition effects through resource demand coefficients. These coefficients lead to non-regulatory interactions among genes that reshape circuit’s behavior. For the activation cascade, such interactions result in surprising biphasic or monotonically decreasing responses. Finally, we use resource demand coefficients to guide the choice of ribosome binding site (RBS) and DNA copy number to restore the cascade’s intended monotonically increasing response. Our results demonstrate how unintended circuit’s behavior arises from resource competition and provide a model-guided methodology to minimize the resulting effects.

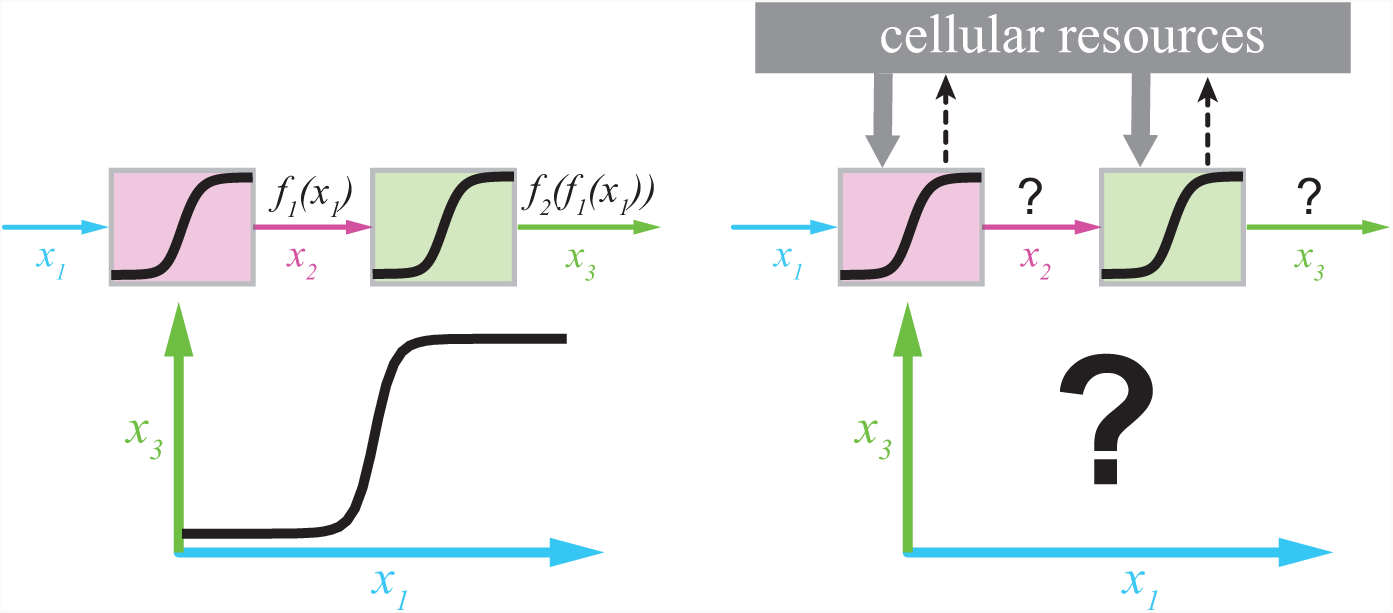

## Introduction

Predicting the behavior of genetic circuits in living cells is a recurring challenge in synthetic biology [1]. Genetic circuits are often viewed as interconnections of gene expression cassettes, which we call *nodes.* Each cassette (node) is composed of core gene expression processes, chiefly transcription and translation. Here, we view each node as an input/output system that takes transcription factors (TFs) as input and gives a TF as output. The input TFs regulate the production of the output TF. Although in an ideal scenario we would like to predict the behavior of a circuit from that of its composing nodes characterized in isolation, in reality, a node’s behavior often depends on its context, including other nodes in the same circuit and the host cell environment [2]. This fact significantly limits our current ability to design genetic circuits that behave as intended. There are a number of causes to context dependence, including unknown structural interactions between adjacent genetic sequences [3], loading of TFs by target DNA sites (retroactivity) [4, 5, 6], unintended coupling between synthetic genes and host cell growth (host-circuit interaction) [7, 8, 9], and competition among synthetic genes with each other for common transcriptional and translational resources [10, 11, 12, 13, 14]. Context dependence due to structural interactions and retroactivity has been addressed by engineering insulation parts and devices [15, 16, 6, 17, 18] and that due to host-circuit interaction may be mitigated to some extent by orthogonal RNA polymerases (RNAPs) and ribosomes [19, 20, 21, 22]. By contrast, the characterization and mitigation of competition for shared resources among synthetic genes remain largely unexplored.

Expression of all genes in a genetic circuit relies on a common pool of transcriptional and translational resources. In particular, the availability of RNAPs and ribosomes has been identified as a major bottleneck for gene expression in bacteria [23, 24, 25, 26]. When a node is activated, it depletes the pool of free RNAPs and ribosomes, reducing their availability to other nodes in the circuit. This can potentially affect the behavior of a circuit altogether. Recent experimental results have demonstrated that competition for transcriptional and translational resources can couple the expression of two synthetic genes that are otherwise unconnected [10, 12]. In particular, limitation in ribosome availability has been identified as the key player in this coupling phenomenon [12]. These works further demonstrate that upon induction of a synthetic gene, the expression level of a constitutively expressed gene on the same plasmid can be reduced by more than 60%. Similar trade-offs have been observed in cell-free systems [27] and in computational models [11, 13, 14, 28].

In this paper, we seek to determine how competition for RNAPs and ribosomes by the genes constituting a synthetic genetic circuit changes the intended circuit’s behavior in bacteria *E. coli.* To address this question, we perform a combined modeling and experimental study. In particular, we develop a general mathematical model that explicitly includes competition for RNAPs and ribosomes in Hill-function models of gene expression. In our models, resource demand coefficients quantify the demand for resources by each node and shape the emergent dose response curve of a genetic circuit. We construct a library of synthetic genetic activation cascades in which we tune the resource demand coefficients by changing the RBS strength of the cascade’s genes and DNA copy number. When the resource demand coefficients are large, the dose response curve of the cascade can either be biphasic or monotonically decreasing. When we decrease the resource demand coefficients, we restore the intended cascade’s monotonically increasing dose response curve. For general circuits, our model reveals that due to non-zero resource demand coefficients, resource competition gives rise to non-regulatory interactions among nodes. We give a general rule for drawing the effective interaction graph of any genetic circuit that combines both regulatory and non-regulatory interactions.

## Results

### Surprising biphasic response of a genetic activation cascade

We built a genetic activation cascade composed of three nodes and two transcriptional activation stages. Two inducer-responsive TFs, LuxR from *Vibrio fischeri* [29] and NahR from *Pseudomonas putida* [30], activate gene expression in their active forms (i.e., *holo* forms) when their respective inducers *N*-hexanoyl-L-homoserine lactone (AHL) and salicylate (SAL) are present. Node 1 uses the *lac* promoter to constitutively express LuxR in a LacI-deficient host strain. By increasing inducer AHL concentration, the active form of LuxR increases, and it can transcriptionally activate the following node. We consider active LuxR as the output of node 1. Node 2 uses transcriptional activation by the active LuxR through the *lux* promoter (Figure 1A). To characterize the dose response curve of this node, we placed red fluorescent protein (RFP) under the control of the *lux* promoter. An increase in AHL concentration increases the active LuxR to promote the production of RFP (Figure 1A).

**Figure 1.**
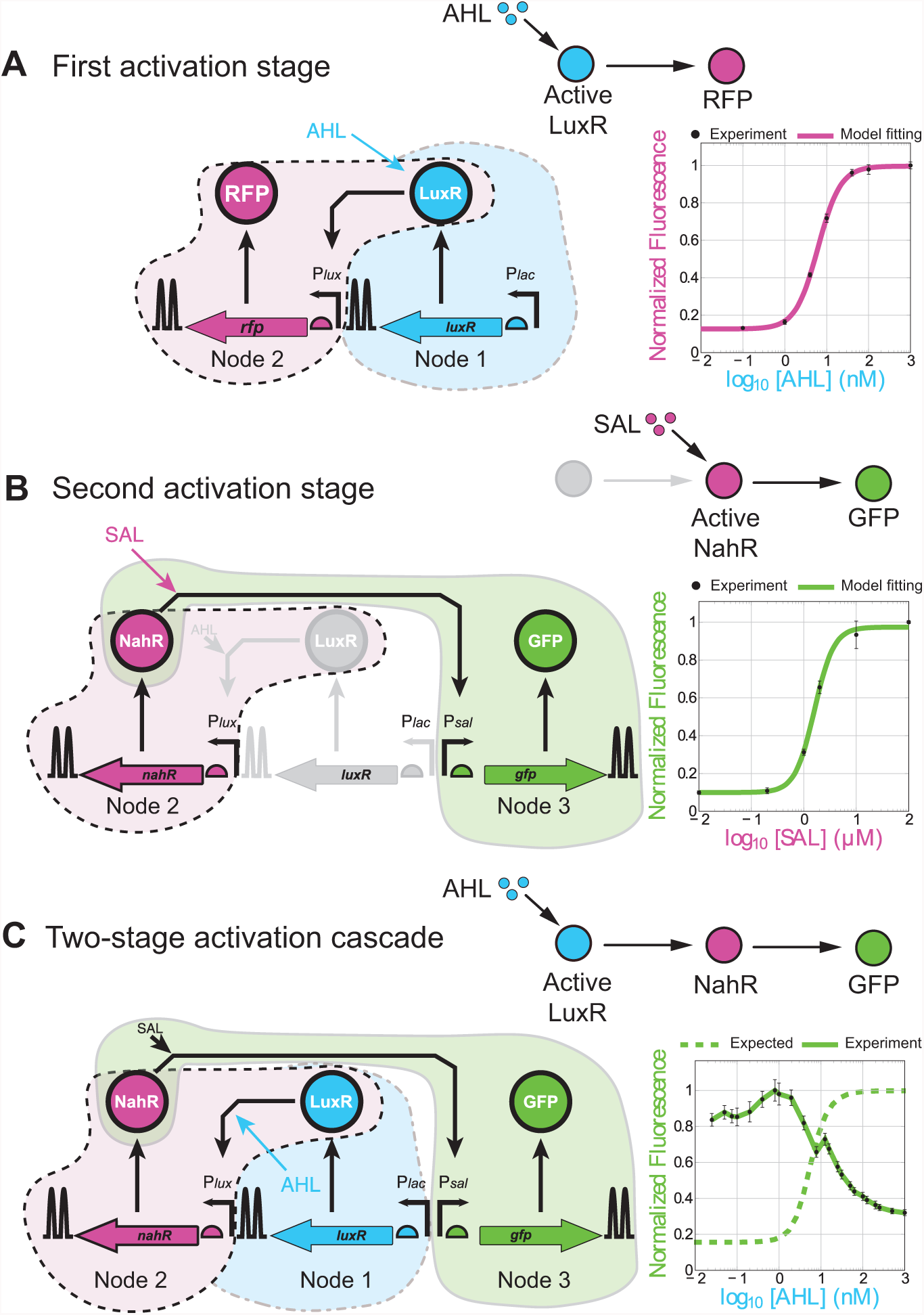
Failure of modular composition in a simple two-stage activation cascade. (A) The first activation stage consists of a node that takes as input the transcription activator LuxR to promote production of RFP as output in the presence of AHL, resulting in a monotonically increasing dose response curve. Upward arrows with leftward/rightward tips represent promoters, semicircles represent RBS, and double hairpins represent terminators. The illustrative diagram composed of nodes and edges at the upper-right corner represents regulatory interactions among species. (B) The second activation stage consists of a node that takes as input the transcription activator NahR to promote production of GFP as output in the presence of SAL, resulting in a monotonically increasing dose response curve. (C) The two-stage activation cascade CAS 1/30 was built by connecting the nodes in a cascade topology. Biphasic dose response curve (solid line) of the cascade was observed instead of the expected monotonically increasing dose response curve (dashed line), which is the composition of the two increasing Hill functions for the individual nodes according to equation (3). All experimental data represent mean values and standard deviations of populations in the steady state analyzed by flow cytometry in three independent experiments. Each plot is normalized to its maximum fluorescence value (see SI Section A7 for details).

Node 3 employs transcriptional activation by active NahR and the *sal* promoter to express green fluorescent protein (GFP) as fluorescence output. Inactive NahR is first produced under the control of the *lux* promoter. We applied a saturating amount of AHL (100 nM) and expressed LuxR constitutively to produce a saturating amount of inactive NahR. By increasing the amount of inducer SAL, active NahR concentration increases, activating production of GFP (Figure 1B). We further confirm that the dose response curve of GFP activation by active NahR remains monotonically increasing under different AHL concentrations (see SI Section A6 and Figure S1).

To build a two-stage activation cascade (CAS 1/30), we connected the three nodes by replacing the RFP in node 2 by NahR. Active NahR can be regarded as the output of node 2 and the input to node 3. With a constant amount of SAL (1 mM), increased AHL concentration leads to increased active LuxR, and hence to increased concentration of active NahR, resulting in increased production of GFP (cascade output).

The expected behavior of this cascade is therefore a monotonically increasing GFP fluorescence as AHL is increased. This can be easily predicted by a standard Hill-function model of the circuit. Specifically, letting Ii denote inducer AHL, I_2_ denote inducer SAL, x_1_ denote active LuxR, x_2_ denote active NahR, and x_3_ denote GFP output, and using *I*_1_*, I*_2_*, x*_1_*, x*_2_ and *x*_3_ (*italics*) to represent their concentrations, we obtain the following ordinary differential equation (ODE) model (see SI Section B1 for details):

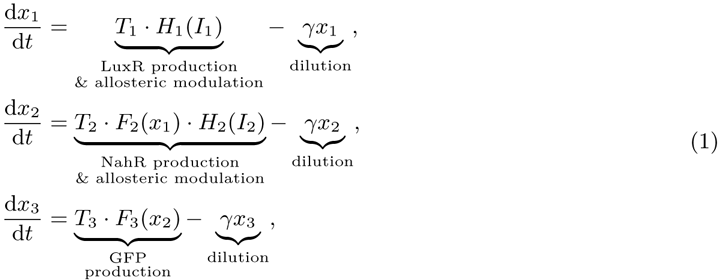

where *T_i_* (*i* = 1, 2, 3) is a lumped parameter describing maximal production rate of node *i* (defined in equation (S15) in SI), *γ* is the dilution rate constant, and *H*_1_(*I*_1_)*, H*_2_(*I*_2_)*, F*_2_(*x*_1_) and *F*_3_(*x*_2_) are standard increasing Hill-functions whose maxima are normalized to 1. *H*_1_(*I*_1_) and *H*_2_(*I*_2_) describe allosteric modulation of LuxR by AHL, and of NahR by SAL. Since we applied a constant amount of SAL (i.e., *I*_2_ = constant), *H*_2_ (*I*_2_) is a constant. Without loss of generality, we assume *H*_2_(*I*_2_) ≡ 1 in sequel, as any non-unity *H*_2_(*I*_2_) can be absorbed into the lumped parameter *T_2_* without affecting our analysis (see SI Section B1). Hill functions *F*_2_(*x*_1_) and *F*_3_(*x*_2_) describe transcriptional regulations of NahR by active LuxR, and of GFP by active NahR, respectively. These Hill functions describing regulatory interactions are derived from the chemical reactions in SI Section B1, and are given by:

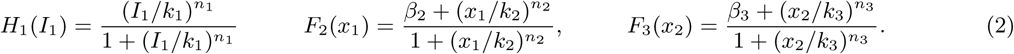

In equations (2), *k_i_* are the dissociation constants between the regulators, I_1_, x_1_ and x_2_, and their respective DNA/protein targets. Dimensionless parameters *β_i_* < 1 characterize basal expressions, and *n_i_* are Hill coefficients capturing cooperativities of the TF and promoter (or of the inducer and TF) bindings. Setting the time derivatives in (1) to zero, we obtain the dose response curve of the cascade as the composition of three increasing Hill-functions:

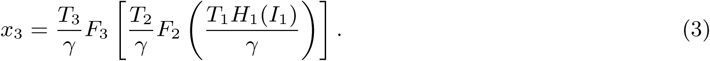

It is clear from (3), that independent of parameters, the steady state of *x*_3_ (GFP concentration) always increases with *I*_1_ (AHL concentration).

Surprisingly, the experimental results contradict this rather trivial prediction. In fact, although the input/output responses of both transcriptionally regulated nodes are monotonically increasing (Figure 1A-B), their cascade shows a biphasic dose response curve, in which the GFP fluorescence decreases with increased concentrations of AHL for higher AHL concentrations (Figure 1C). This fact clearly demonstrates that while the standard model well represents the activation behavior of each individual node, its predictive ability is lost when the nodes are connected and thus are simultaneously activated.

In the next section, we derive a new model, similar in form to that of model (1), which is able to predict the experimentally observed behavior.

## A cascade model taking into account resource competition predicts nonregulatory interactions

An underlying assumption in the standard Hill-function model (1) is that the concentrations of free RNAPs and ribosomes can be regarded as constant parameters [31, 32] (refer to SI Section B1). In reality, because their total availability is limited [23, 24, 26], their free concentrations should depend on the extent to which different nodes in a circuit demand them. With reference to Figure 1C, the biphasic response of *x*_3_ can be explained by the following resource competition mechanism. When we increase *I*_1_., the concentration of x_1_ increases, promoting production of x_2_. As node 2 sequesters more RNAPs and ribosomes, the amount of free resources decreases, which in turn result in decreased transcription and translation rates in node 3.

We therefore created a model that explicitly accounts for the limited concentrations of RNAPs and ribosomes and for their competition by the three nodes in the cascade. For a given growth rate, the total concentrations of RNAPs and ribosomes can be assumed constant parameters [9, 23]. Considering the conservation law for these resources and solving for their free concentrations (see SI Section B2), we obtain the following modified Hill-function model:

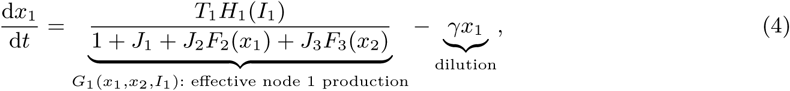

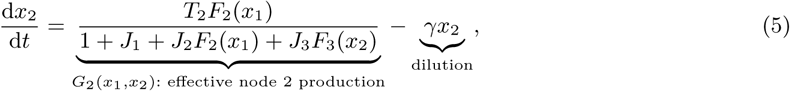

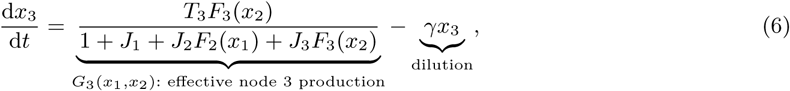

in which *H*_1_(*I*_1_)*, F*_2_(*x_1_*) and *F*_3_(*x*_2_) are regulatory Hill functions defined in equations (2), and *T_i_* (*i* = 1, 2, 3) is the maximum production rate of node *i* defined in (S25) in SI. The lumped dimensionless parameter *J_i_* can be understood as an indicator of maximal resource demand by node i, and we call it *resource demand coefficient*. It is defined as:

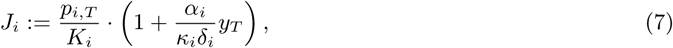

where *p_i,T_* is the DNA copy number of node *i*; *α_i_* is its transcription elongation rate constant, describing the average number of mRNAs transcribed from a single DNA molecule in unit time; *δ_i_* is mRNA decay rate constant, and *y_T_* is the total concentration of RNAPs. The ability of each DNA molecule (mRNA molecule) to occupy free RNAPs (ribosomes) is characterized by lumped coefficient *K_i_* (*κ_i_*), defined in equations (S3) and (S8) in SI. They can be viewed as effective dissociation constants that decrease with (i) stronger affinity between activated promoter (RBS) in node *i* and free RNAPs (ribosomes), and (ii) lower transcription (translation) elongation rate constants. Physically, resource demand coefficient of node *i* (*J_i_*) increases as (I) the total number of promoter sites (*p_i_,_T_*) increases, (II) the total number of mRNA molecules (*p_i_*,*_T_* α*_i_y_T_*/*K_i_δ_i_*) increases, (III) the ability of each DNA molecule to sequester free RNAPs (1/*K_i_*) increases, or (IV) the ability of each mRNA molecule to sequester free ribosomes (1/ *κ_i_*) increases. For a given transcriptional activation level, the portion of resources allocated to each node is quantified by J_1_, *J*_2_*F*_2_(*x*_1_) and *J*_3_*F*_3_(*x*_2_), respectively, and follow the conservation law (see SI Section B5 for derivation):

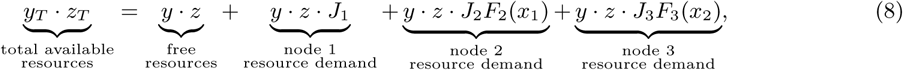

where *y_T_* (*y*) and *z_T_* (*z*) are the total (free) amount of RNAPs and ribosomes, respectively.

The major difference between model (4)-(6) and the standard Hill-function model (1) is the common denominator 1 + *J*_1_ + *J*_2_*F*_2_(*x*_1_) + *J*_3_*F*_3_(*x*_2_) in the effective node production rates *G*_1_(*x*_1_,*x*_2_*,I*_1_), *G*_2_(*x*_1_*,x*_2_) and *G*_3_(*x*_1_,*x*_2_). In a resource-abundant situation where RNAPs and ribosomes bound to all nodes are much smaller than their free amounts (*y* ≈ *y_T_* and *z* ≈ *z_T_*), we have *J*_1_*,J*_2_*,J*_3_ ≪ 1, and model (4)-(6) reduces to the standard Hill-function model (1). Detailed proof of this result is in SI Section B5. Because of the common denominator, the production of each node depends on all TFs present in the circuit as opposed to depending only on its own inputs as in equation (1). In particular, regardless of parameters, we always have the following effective interactions among the cascade nodes (see SI Section B3.1 for derivation):

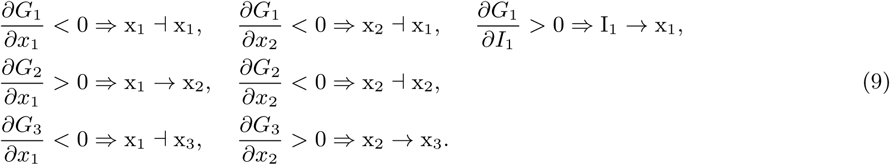

While interactions I_1_ → x_1_, x_1_ → x_2_ and x_2_ → x_3_ are due to the intended allosteric modulation and transcriptional activations, the other interactions are not present in the standard model (1). They can be regarded as non-regulatory interactions arising from resource competition among nodes. In particular, the non-regulatory interactions x_1_ ⊣ x_3_ and x_1_ ⊣ x_1_ are due to the fact that as *x*_1_ increases, production of x_2_ is activated, depleting the pool of free resources, thus reducing the amount of resources available to initiate transcription and translation of x_3_ and x_1_, respectively. Similarly, an increase in *x*_2_ activates production of x_3_, reducing resources available to its own expression and that of x_1_, leading to non-regulatory interactions x_2_ ⊣ x_2_ and x_2_ ⊣ x_1_.

Based on (9, the effective interactions among nodes in an activation cascade are shown in Figure 2A, where we use red dashed edges to represent emergent non-regulatory interactions due to resource competition. The non-regulatory interactions create a feed-forward edge x_1_ ⊣ x_3_, a feedback edge x_2_ ⊣ x_1_ and two negative auto-regulation edges: x_1_ ⊣ x_1_, x_2_ ⊣ x_2_. In SI Section B3.1, we demonstrate that regardless of emergent negative auto-regulation edges on x_1_ and x_2_, and the feedback edge x_2_ ⊣ x_1_, *x*_1_ and *x*_2_ still increases with inducer input *I*_1_ as expected. Therefore, the topology of this activation cascade effectively becomes a type 3 incoherent feed-forward loop (IFFL) [31], where x_3_ production is jointly affected by regulatory activation from x_2_ and non-regulatory repression from x_1_. It is well-known that, depending on parameters, the dose response curve of an IFFL can be monotonically increasing, decreasing or biphasic [33, 34]. As we increase *I*_1_ to increase *x*_1_, if transcriptional activation x_1_ → x_2_ → x_3_ is stronger than non-regulatory repression x_1_ ⊣ x_3_, then the dose response curve is monotonically increasing. Conversely, if the non-regulatory repression is stronger than transcriptional activation, the dose response curve becomes monotonically decreasing. Biphasic responses can be expected when transcriptional activation dominates at lower inducer levels, and resource-competition-induced non-regulatory repression becomes more significant at higher inducer levels. A detailed analytical treatment is in SI Section B3.

**Figure 2.**
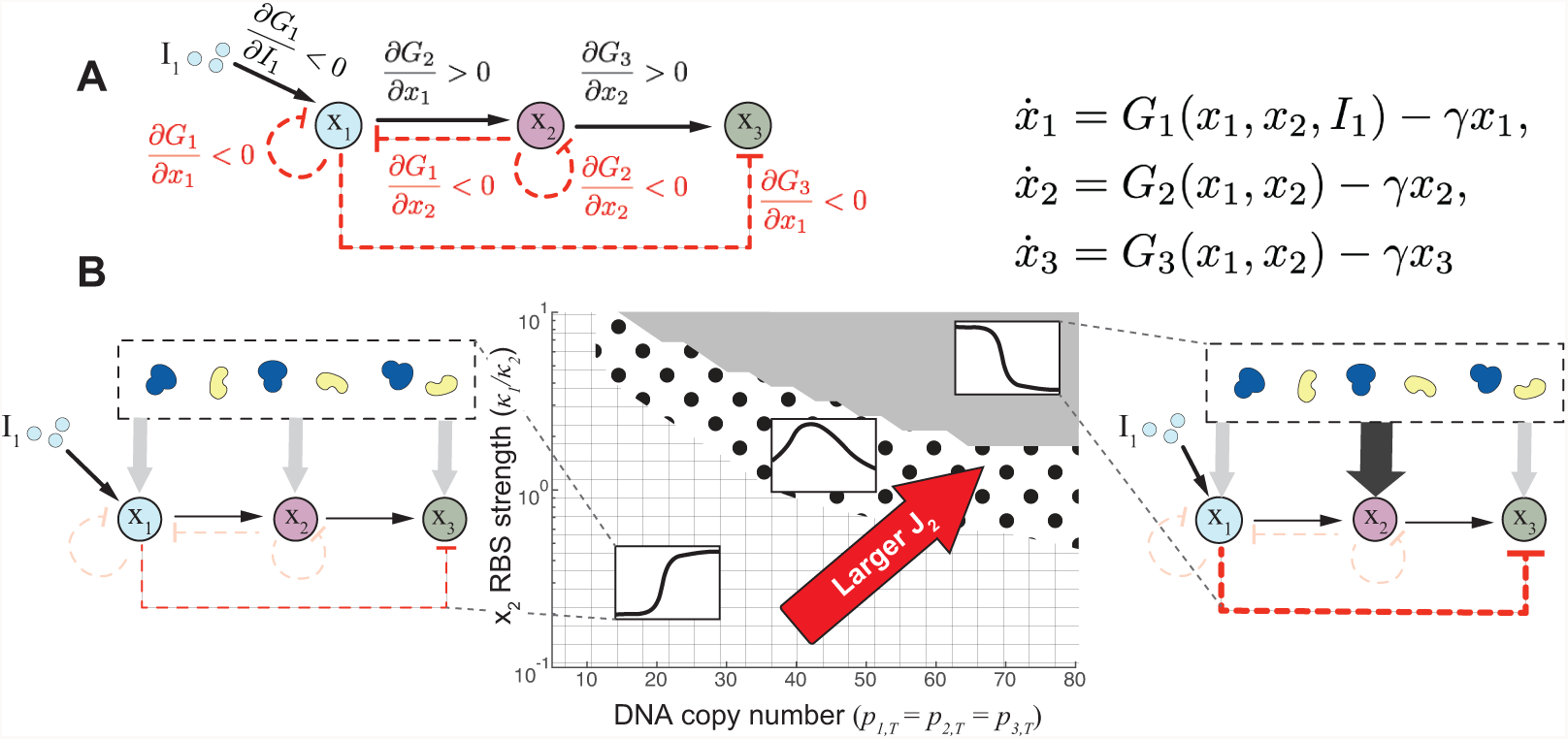
Activation cascade becomes an effective type 3 **IFFL** due to resource competition. (A) Effective interactions among nodes in a two-stage activation cascade with resource competition. Black solid edges are regulatory interactions, and red dashed edges represent emergent non-regulatory interactions due to resource competition. (B) Parameter space illustrating diverse cascade dose response curves obtained from numerical simulations when the resource demand coefficient *J*_2_ is changed. The horizontal axis shows the DNA copy number, and the vertical axis shows the RBS strength of node 2. Numerical values on the vertical axis represent the ratio between the dissociation constant of node 1 (*κ*_1_) between RBS and ribosomes (kept constant at 15 *μ*M), and that of node 2 (*κ*2). The cascade has monotonically decreasing, biphasic or monotonically increasing dose response curve depending on whether the parameters fall into the gray, dotted or grid shaded region in the parameter space, respectively. Simulations are based on a full reaction rate equation model corresponding to the chemical reactions in SI Section B1 and B2. Parameter values are listed in SI.

The strength of the non-regulatory repression x_1_ ⊣ x_3_ can be reduced by decreasing resource demand coefficient of node 2 (*J*_2_). This is because, as a result, the dose response curve of an activation cascade is monotonically increasing when *J*_2_ ≪ 1 (see SI Section B3.2). Conversely, we expect the dose response curve to be monotonically decreasing when *J*_2_ is large, and to be biphasic for intermediate values of *J*_2_. Based on the definition of resource demand coefficient in (7), we can decrease *J*_2_ by choosing weak node 2 RBS strength and low DNA copy number. We simulated the dose response curves of activation cascades with different node 2 RBS strengths and DNA copy numbers, presented in the parameter space in Figure 2B. The lower left corner of the parameter space corresponds to the cascade with the smallest *J*_2_, and the upper right corner corresponds to the largest *J*_2_. In accordance with these predictions, simulations in Figure 2B confirms that smaller *J_2_* (weak x_2_ RBS and low DNA copy number) results in monotonically increasing response (grid shaded region), while larger *J_2_* (strong x_2_ RBS and high DNA copy number) results in monotonically decreasing response (gray region). The dotted region corresponds to intermediate values of *J*_2_ which result in biphasic response.

## Model-guided design recovers monotonically increasing response of the cascade

Based on the simulation map in Figure 2B and the mathematical analysis of model (4)-(6) described in the previous section, we created a library of activation cascades in which each cascade should result into one of the three different behaviors shown in Figure 2B. This library is composed of cascades that differ in the value of the resource demand coefficient of NahR (*J*_2_), with the rationale that we can mitigate the strength of the key non-regulatory interaction x_1_ ⊣ x_3_ to recover the intended monotonically increasing dose response curve of the cascade. In particular, starting from CAS 1/30, whose dose response curve is biphasic (Figure 3A), we designed circuit CAS 0.3/30 with about 30% RBS strength [12] of NahR compared to CAS 1/30, theoretically resulting in a reduction of *J*_2_. We therefore expect a reduction of the x_1_ ⊣ x_3_ interaction strength, leading to a monotonically increasing dose response curve, which is confirmed by the experiment (Figure 3B).

**Figure 3.**
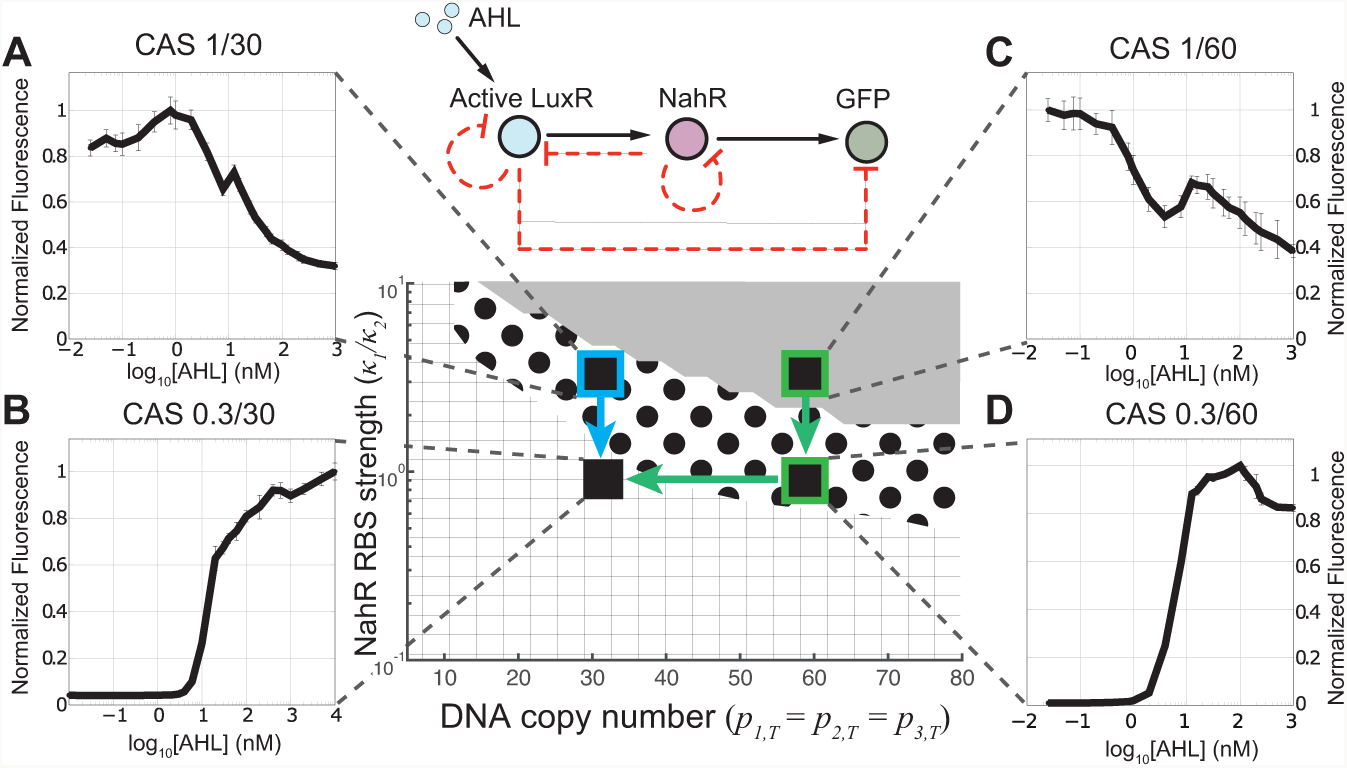
Model-guided design restores the monotonically increasing dose response curve of the cascade. The dose response curves of circuits CAS 1/30 (A) and CAS 1/60 (C) were biphasic and monotonically decreasing, respectively. By reducing the RBS strength of NahR, the dose response curve of CAS 0.3/30 (B) became monotonically increasing, and that of CAS 0.3/60 (D) became biphasic. Further decreasing the copy number of CAS 0.3/60 to CAS 0.3/30 restored the monotonically increasing dose response curve. Experimental results are presented on top of the parameter space created in Figure 2B by simulations. Blue and green arrows represent design actions to restore the monotonically increasing dose response curves starting from failed cascades CAS 1/30 and CAS 1/60, respectively. Mean values and standard deviations of fluorescence intensities at the steady state are calculated from three independent experiments analyzed by flow cytometry and normalized to the maximum value in each plot (see SI Section A7).

Similarly, we constructed another cascade circuit CAS 1/60 in which the DNA copy number is about twice as that of CAS 1/30 (about 60 vs 30). According to our model, resource demand coefficient of NahR *J*_2_ in CAS 1/60 should double compared to that of circuit CAS 1/30. Therefore, we expect a possibly monotonically decreasing dose response curve. Experiments confirm this prediction (Figure 4C). A local increase in GFP fluorescence at about 10 nM AHL is due to the two-step multimerization of NahR proteins [35], which is detailed in SI Section A5. To obtain a monotonically increasing dose response curve from this circuit, we first reduced NahR resource demand coefficient *J*_2_ by designing a circuit CAS 0.3/60, whose NahR RBS strength is 30% compared to that of CAS 1/60. Theoretically, depending on parameters, reduced *J*_2_ can lead to either monotonically increasing or biphasic dose response curves (see Figure 2B). Our experiment show that the response of CAS 0.3/60 is indeed biphasic (Figure 3D). To restore a monotonically increasing dose response curve, we can further decrease *J*_2_ by reducing DNA copy number to create circuit CAS 0.3/30, whose dose response curve is monotonically increasing (Figure 3B).

**Figure 4.**
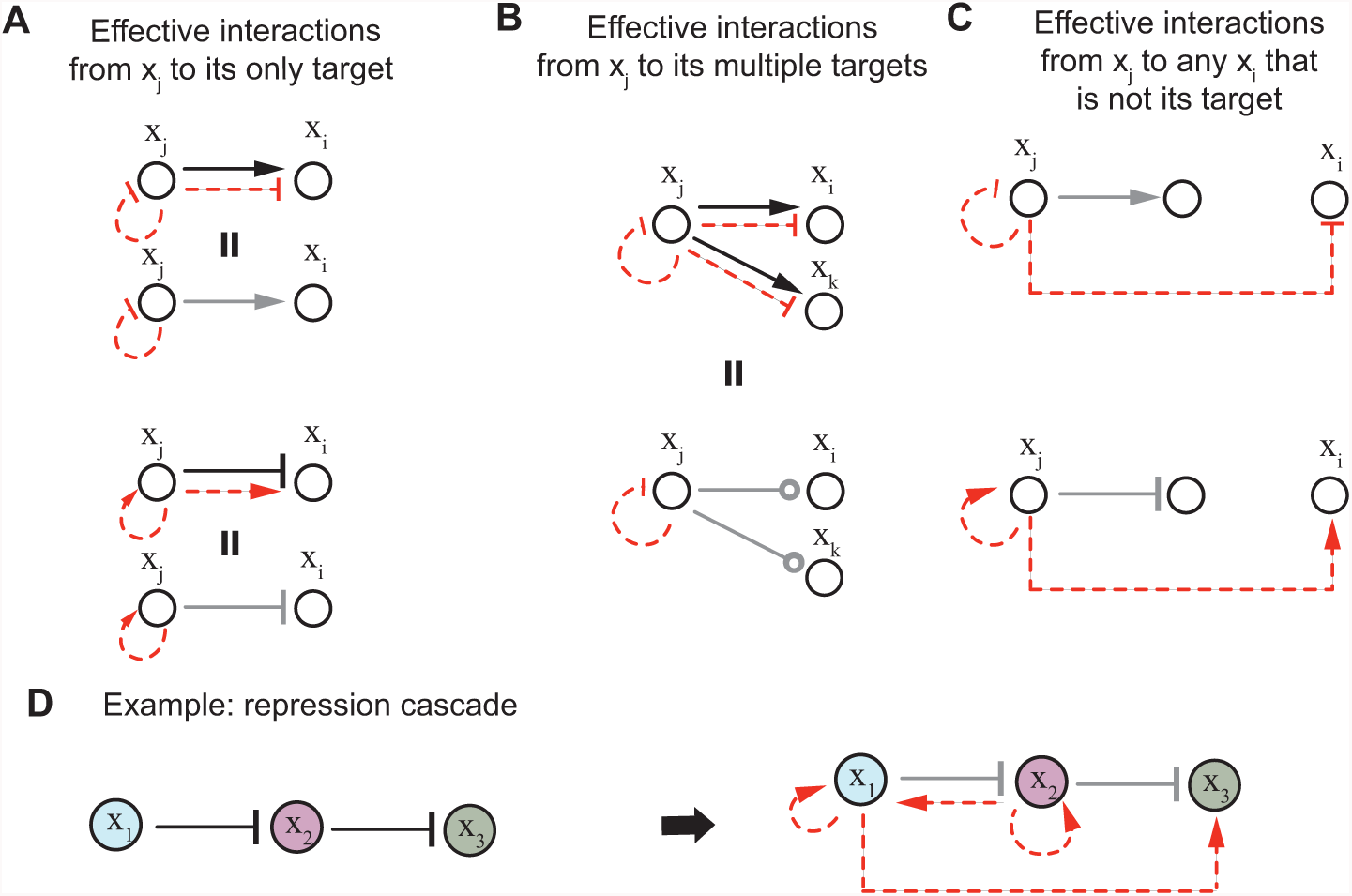
Rules to determine effective interaction graphs arising from resource competition in a genetic circuit. Black solid edges represent regulatory interactions; red dashed edges represent nonregulatory interactions arising from resource competition; if a black and a red edge have the same starting and ending nodes, we indicate their combined effect with a gray edge. (A) If TF x_j_ has only one target, then resource competition does not change the nature (activation or repression) of interaction from x_j_ to its target. However, it weakens the intended strength. (B) If TF x_j_ regulates multiple targets, then the effective interactions from x_j_ to its targets are undetermined. (C) If x_j_ is a transcriptional activator (repressor), then it becomes an effective repressor (activator) for all nodes that are not its target. (D) Applying the rules in A-C, we determine the effective interaction graph for a repression cascade.

## General rules to draw effective interactions in genetic circuits

Interaction graphs, which use directed edges to represent regulatory interactions, are a convenient graphical tool to design and/or analyze the qualitative behavior of a genetic circuit [31]. Here, we expand the concept of interaction graph to incorporate non-regulatory interactions due to resource competition. We call the resultant interaction graph an *effective interaction graph*, which includes both regulatory interactions and non-regulatory interactions due to resource competition. In an effective interaction graph, we draw x → y (x ⊣ y) to represent effective activation (repression). We draw x ⊸ y if the interaction is undetermined, that is, it depends on parameters and/or x concentration.

The resource competition model (4)-(6) and the effective interaction graph identified in Figure 2A for the activation cascade can be generalized to any genetic circuit in a resource-limited environment. Each node *i* is a system that takes active TFs as inputs through the process of transcriptional regulation, and produce an active TF x_i_ as an output. Therefore, each node represents a dynamical process that can be captured by the ODE describing the rate of change of active xi’s concentration *x_i_.* In a circuit with *N* nodes, we can write the dynamics of node *i* as (see SI Section B4 for derivation):

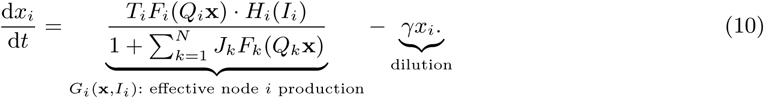

According to model (10), effective production rate of *x_i_, G_i_*(x*,I_i_*), is jointly affected by transcriptional regulation *T_i_F_i_*(*Q_i_*x), allosteric modulation *H_i_*(*I_i_*), and resource competition *R*(x):= 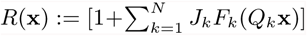.

In particular, since resources are shared among all nodes, *R*(x) is a common denominator to the effective production rate of every node. In model (10), *T_i_* and *J_i_* are lumped parameters that represent the maximum production rate of x_i_ and the maximum resource demand by node *i*, respectively (see equation (S70) and (7) for precise expressions). In particular, we gave the expression of *J_i_* in equation (7) along with the explanation of its physical meaning. The binary matrix *Q_i_* selects the TF inputs to node *i* (see SI Section B4.1 for precise definition), and the vector x = [*x*_1_,…*,x_N_*]^*T*^ represents the concatenation of all active TFs’ concentrations in the circuit. Active TFs are defined as proteins, either in inducer bound or unbound form, that can transcriptionally regulate a gene. Normalized Hill function *F_i_*(*Q_i_*x) describes the transcriptional regulation of node *i* by its input TFs (see SI equation (S53) for precise expression). For those TFs whose activity can be allosterically modulated by an inducer I_i_, normalized Hill function *H_i_* (*I*_*i*_) represents the portion of TF that is active. If node *i* is not transcriptionally regulated, that is, x_i_ is constitutively expressed, then *F_i_* ≡ 1. Similarly, if the activity of the TF x_i_, is not allosterically modulated by an inducer, then *H_i_* ≡ 1. Parameter *γ* is a dilution rate constant that models cell growth.

Given model (10), a *regulatory interaction* from x_j_ to x_i_ is given by a non-zero ∂*F_i_*/*∂x_j_*. It is an activation (→) if *∂F_i_*/*∂x_j_*> 0 and a repression (⊣) if *∂F_i_*/*∂x_j_* < 0. A *non-regulatory interaction* due to resource competition from xj to any node x_i_ is present if *∂R*/*∂x_j_* is non-zero. Since *R*(x) is in the denominator, conversely, the non-regulatory interaction is an activation (→) if *∂R/∂x_j_* < 0 and a repression (⊣) if *∂R/∂x_j_*> 0. The quantity *∂R/∂x_j_* captures the following physical phenomenon, which is responsible for non-regulatory interactions. As the concentration of active TF x_j_ increases, resource demand by the nodes that x_j_ activates/represses, which we call x_j_’s targets, increases/decreases; this in turn, reduces/increases the free amount of resources available to all nodes in the circuit. Therefore, the existence of a non-regulatory interaction originating from node *j* is exclusively dictated by the action (activation or repression) that TF x_j_ exerts on its targets; it is not dictated by the pure change in the concentration of TF x_j_ itself. *Effective interaction* from x_j_ to x_i_ represents the combined effect of regulatory and non-regulatory interaction from x_j_ to x_i_, and is identified based on the sign of *∂G_i_/∂x_j_*.

Following the above, we list a set of immediate graphical rules to draw the effective interactions originating from x_j_ based on whether it is a transcriptional activator or repressor and based on the number of its targets (Figure 4A-C). These rules establish that when x_j_ transcriptionally regulates only one target, the nature of the effective interaction (i.e. activation vs. repression) from x_j_ to its target is unaffected by resource competition, but the strength of such interaction is weaker than the intended regulatory interaction. (Figure 4A). However, when x_j_ has multiple targets, the nature of effective interactions from x_j_ to its targets are undetermined (see Figure 4B and example in Figure S11). If x_j_ is a transcriptional activator (or repressor), then it is effectively repressing (or activating) all nodes that are not its targets, possibly including itself (Figure 4C). Detailed derivation of these graphical rules can be found in SI Section B6. Using these rules, the effective interaction graph of a two-stage activation cascade (Figure 2A) can be immediately identified. In Figure 4, we use black solid edges to represent regulatory interactions, and red dashed edges to represent non-regulatory interactions due to resource competition. If a black and a red edge have the same starting and ending nodes, we indicate their combined effect with a gray edge.

As an additional example of these graphical rules, we construct the effective interaction graph of a two-stage repression cascade in Figure 4D. Both x_l_ and x_2_ are repressors with only one target. Therefore, applying the rule in Figure 4A, we obtain x_l_ ⊣ x_2_ and x_2_ ⊣x_3_. Since x_3_ is not a target of x_l_ and x_l_ is a repressor, applying the rule in Figure 4C, we obtain x_l_ → x_3_. Similarly, x_l_ and x_2_ are effectively activating themselves, and x_2_ effectively activates x_l_. Since x_3_ does not transcriptionally activate or repress a target, there is no effective interaction originating from x_3_. The resultant effective interaction graph in Figure 4D leads to a dose response curve that is monotonically increasing regardless of parameters (refer to SI Section B7). We can further use these interaction graphs to compare circuits with same functionality. Specifically, with a positive inducer input, the activation cascade of Figure 2A and the repression cascade of Figure 4D both are intended to have a monotonically increasing dose response curve. Since the repression cascade can keep this qualitative behavior in the face of resource competition, while the activation cascade may not, the former design is more robust to resource competition than the latter.

## Discussion

Gene expression relies on transcriptional and translational resources, chiefly RNAPs and ribosomes. As all genes in a circuit compete for these limited resources, unintended non-regulatory interactions among genes arise. These interactions can dramatically change the intended behavior of a genetic circuit. In this paper, through a combined modeling and experimental study, we have characterized the extent to which resource competition affects a genetic circuit’s behavior. We have incorporated resource competition into standard Hill-function models through resource demand coefficients, which can be readily tuned by key circuit parameters such as RBS strength and DNA copy number. These coefficients dictate the strengths of non-regulatory interactions and can be effectively used to guide the design of a genetic circuit toward the intended behavior. Our mathematical model further provides a simple graphical tool to identify the nature of non-regulatory interactions (i.e. activation vs. repression) and to create the effective interaction graph of the circuit. Under the guidance of the model, we created a library of genetic activation cascades and demonstrated that, by tuning the resource demand coefficients of the cascade’s nodes, the strengths of non-regulatory interactions can be predictably controlled and intended cascade’s response can be restored.

Previous theoretical studies have analyzed how competition for shared resources affects gene expression. Using a stochastic model [13], Mather et al. found a strong anti-correlation of the proteins produced by ribosome-competing mRNAs. Rondelez [28] developed a general model to describe substrates competing for a limited pool of enzymes. De Vos et al. [11] analyzed the response of network flux toward changes in total competitors and common targets. More recently, Raveh et al. [14] developed a ribosome flow model to capture simultaneous mRNA translation and competition for a common pool of ribosomes. In [12], Gyorgy et al. developed a mechanistic resource competition model that gives rise to “isocost lines” describing tradeoffs in gene expression, which were experimentally validated. All these models, with the exception of [28], are restricted to circuits without regulatory links among competing nodes. In contrast, our general model explicitly accounts for regulatory interactions among nodes and reproduce the “isocost lines” of [12] as a special case (see SI Section B4.4). Furthermore, differently from [28], our model couples the resources’ enzymatic reactions with the slower gene expression reactions to obtain a model for resource-limited genetic circuits.

Previous experimental studies have provided evidence that transcriptional and translational resources may be limited in the cell by showing that DNA copy number, mRNA concentration, and protein concentration do not always linearly correlate with each other [24, 26]. Accordingly, there has been extensive experimental evidence that synthetic genes’ over-expression inhibits host cell growth [7, 36, 8, 9]. However, the effects of competition for shared resources on genetic circuits have only been recently addressed, mostly focusing on the single-gene effects as opposed to investigating the emergent effects at the network level [10, 12, 22, 37]. In this paper, we have theoretically predicted and experimentally demonstrated that significant network-level effects arise due to non-regulatory interactions dictated by resource competition. These interactions need to be accounted for in circuit design and optimization. Accordingly, we have provided a model-based approach to guide genetic circuit design to mitigate the effects of unintended interactions.

As a form of host-circuit interaction, previous studies have shown that overexpression of synthetic genes may retard host cell growth, which in turn affects dilution rates of the synthetic species [7, 8, 9, 38]. In our experiments with CAS 0.3/30 and CAS 0.3/60, none to very modest changes in growth were observed. In experiments with CAS 1/60 and CAS 1/30, appreciable decrease in growth rates were observed when AHL concentrations are higher than about 10nM (see Figure S3). However, unintended effects of resource competition can already be observed in the dose response curve of CAS 0.3/60 for all AHL concentrations and those of CAS 1/30 and CAS 1/60 for low AHL concentrations. Nevertheless, in SI Section B8, we use a simple model of dilution rate constant modulation by resource depletion to demonstrate that the qualitative behavior of the activation cascade model is unchanged even when growth retardation is taken into account.

As circuits grow in size and complexity, a “resource-aware” design approach needs to be adopted by synthetic biologists. While resource competition can be exploited in certain situations to our advantage [39, 40, 41], its global and nonlinear features largely hamper our capability to carry out predictive design. To alleviate the effects of resource competition, metabolic engineers down-regulate undesired gene expression to re-direct resources to the pathway of interest, thus increasing its yield [42, 43]. Similarly, in a genetic circuit, we can tune the resource demand coefficients of nodes by selecting appropriate RBS and DNA copy numbers to diminish the resource demand by certain nodes and hence make more resource available to other nodes. This tuning should be performed by keeping in mind other design specifications that the circuit may have, such as maximal output or sensitivity of the dose response curve [1]. A simulation example of how to relate easily tunable parameters, such as RBS strength and DNA copy number, to circuit’s output is given in SI Section B3.3 for the genetic activation cascade. At the higher abstraction level of circuit topology, our model helps to identify topologies whose behavior is less sensitive to the effects of non-regulatory interactions. We provided an example of this with the two-stage activation and repression cascades. While the dose response curve of the former can be completely reshaped by non-regulatory interactions due to resource competition, the dose response curve of the latter is independent of resource competition.

Characterization of resource competition has deep implications in the field of systems biology, in which a major task is to reconstruct networks from data. In this case, it is critical to distinguish direct regulatory interactions from indirect ones [44], which may arise from non-regulatory interactions due to resource competition. In this sense, our model may provide deeper insights to guide the identification of natural networks from perturbation data.

## Associated content

Methods and materials, detailed experimental data and mathematical models are described in Supporting Information.

## Acknowledgements

We thank Fiona Chandra, Andras Gyorgy and Carlos Barajas for helpful discussions and suggestions. YQ and HHH were supported in part by AFOSR grant FA9550-14-1-0060 and ONR award N000141310074. JJ was supported by BBSRC grant BB/M009769/1.

## Author contributions

All authors designed experiments and interpreted the data; HHH, JJ, YQ performed experiments; HHH and JJ cloned constructs; YQ developed the mathematical model and performed simulations; YQ and HHH wrote the paper; DDV edited the paper, designed the research and assisted with developing the mathematical model.

**References**

